# Polyfunctional pathogen-specific CD4^+^ T cells reside in the lungs and tumors of NSCLC patients

**DOI:** 10.1101/2022.02.16.479923

**Authors:** Anna E. Oja, Florencia Morgana, Ruth Hagen, Cherien A. Ghandour, Martijn A. Nolte, Suzan H.M. Rooijakkers, René A.W. van Lier, Bart W. Bardoel, Pleun Hombrink

## Abstract

Local T cell responses are required for optimal protection of the lungs against airborne pathogens. This localized protection is mediated by various immune cells, including resident memory T cells (T_RM_). While human lung CD8^+^ T_RM_ line the epithelium and are enriched for recognition of respiratory viruses, we found CD4^+^ T_RM_ to exhibit more heterogeneous localization patterns, surrounding airways, forming clusters in the lung parenchyma, and lining the epithelium. This heterogeneity was also reflected functionally, as lung CD4^+^ T_RM_ were enriched for recognition of diverse classes of respiratory pathogens. Upon stimulation, lung CD4^+^ T_RM_ expressed different polyfunctional cytokine profiles depending on the pathogen recognized. CD4^+^ T_RM_ responding to respiratory viruses and bacteria were biased for production of IFN-γ and IL-17, respectively. Strikingly, pathogen-specific CD4^+^ T_RM_ also represent a significant fraction in NSCLC tumors that remained polyfunctional despite high PD-1 expression. These findings are not only important for vaccine design, but also provide a rationale for reinvigorating anti-tumor immunity through triggering of polyfunctional pathogen-specific CD4^+^ tumor infiltrating lymphocytes.

## Introduction

The human lungs are our largest interface with the outside environment. With every breath the pulmonary epithelium is exposed to airborne particles containing various classes of infectious pathogens. Furthermore, evidence of a comprehensive lung microbiota has emerged, and its role in regulating the balance of immune tolerance and activation as well as lung carcinogenesis^1,2^. Pathogens employ different routes of entry and mechanisms to infect and/or colonize the lungs and tailor-made heterotypic immune responses are required to sustain lung homeostasis. However, sometimes the immune system fails to protect against infections of opportunistic, and/or novel pathogens. Consequently, frequently mutating pathogens, such as seasonal influenza strains and the newly mutated severe acute respiratory syndrome coronavirus 2, SARS-CoV-2, remain global threats. T cells are an integral part of local adaptive immune responses and understanding their role in maintaining pulmonary immune homeostasis is crucial for the design of therapeutic interventions, such as vaccines.

Long-term memory T cells remain after the clearance of a pathogen and are designed to prevent secondary infections. Deposition of T cells at former sites of pathogen entry is pivotal to this task. These cells are known as resident memory T cells (T_RM_), as they reside within tissues, in disequilibrium to the circulation, to mediate localized protection^3^. CD8^+^ T_RM_ inhabit various tissues, including the lungs, and their protective roles against secondary lung infections are evident^4,5^. In mice, lung CD8^+^ T_RM_ create niches around sites of regeneration following tissue damage^6^. Human CD103^+^CD8^+^ T_RM_ line the lung epithelium and are programmed for tissue surveillance^7,8^. Furthermore, CD103^+^CD8^+^ T_RM_ are enriched for recognition of localized viruses, influenza and RSV, but not systemic viruses, CMV or EBV^7^. The localization of CD8^+^ T_RM_ is key for rapid responses to secondary pathogen encounter.

The protective capacity of CD4^+^ T_RM_ is not well defined. CD4^+^ T_RM_ reside in both lymphoid and non-lymphoid organs in mouse and man^4,9^. While heterogeneity among CD4^+^ T_RM_ has been described on a phenotypic and transcription level^10,11^, the functional foundation of such heterogeneity is poorly understood. Whereas CD8^+^ T_RM_ mainly contribute to clearance of viral infections, CD4^+^ T_RM_ protect against intracellular viruses and bacteria and extracellular bacteria, and fungi. Adoptive transfer of CD4^+^ T_RM_, but not circulating CD4^+^ T cells, protect naïve mice against lethal influenza infection^12,13^. Furthermore, *Streptococcus pneumoniae* infection induces heterotypic immune protection by lung CD4^+^ T_RM_^14^. On the contrary, in *Aspergillus fumigatus* infection, migratory, but not resident CD4^+^ T cells, provide optimal protection^15^ and in chronic *A. fumigatus* infection CD4^+^ T_RM_ are even detrimental^16^. Thus, lung CD4^+^ T_RM_ can be protective or pathogenic, depending on the context. Polyfunctional Th1- and Th17-type CD4^+^ T_RM_ persist in the human lungs^10,11^, but whether these cells recognize viral, bacterial, or fungal antigens and how they localize within the lungs remains elusive.

T cells also play a central role in immune responses to cancer^17^. Tumors contain tumor-specific CD8^+^ and CD4^+^ tumor infiltrating lymphocytes (TILs), recognizing neoantigens^18^. Furthermore, tumor infiltration by CD8^+^ TILs with a T_RM_ phenotype associate with a favorable prognosis in NSCLC^19^. For the optimal formation of cytotoxic CD8^+^ T cells against tumors, CD4^+^ T cell help is required^20^. However, dysfunctionality or exhaustion of the tumor-specific TILs is a major obstacle in anti-cancer immune therapies. Furthermore, a large proportion of CD8^+^ TILs do not recognize autologous tumor cells^21^. At least a part of these non-tumor-specific CD8^+^ TILs are virus-specific CD8^+^ T cells with a residency phenotype, known as bystander cells. Bystander CD8^+^ TILs infiltrate various solid tumors, including lung tumors^22,23^. NSCLC tumors also contain substantial frequencies of non-regulatory CD4^+^ TILs with a residency phenotype^24^. However, whether these TILs contain pathogen-specific CD4^+^ T_RM_ is not known.

Here, we show that CD4^+^ T_RM_ localize in different patterns throughout the human lungs. This heterogeneous spatial organization was also reflected by functional diversity of CD4^+^ T_RM_. Polyfunctional CD4^+^ T cells recognizing viral, bacterial, and fungal antigens with distinct cytokine profiles are enriched in the lungs and tumors of NSCLC patients compared to peripheral blood. Our results offer a first glance into what drives CD4^+^ T_RM_ heterogeneity in the human lungs. Further studies are necessary to determine how to exploit these cells for heterotypic protection of vaccines and reinvigorating the tumor microenvironment.

## Results

### Human lung CD4^+^ T_RM_ display heterogeneous spatial organization patterns

Pathogen-specific CD4^+^ T_RM_ should be strategically localized within the lung to respond quickly and vigilantly upon pathogen entry. As various classes of airway pathogens exploit different mechanisms of entry, we hypothesized that this may be reflected by the spatial organization of different CD4^+^ T_RM_ subsets. CD69 and CD103 are often used to identify T_RM_^8,10^. Therefore, we investigated the localization of CD103^+^ and CD69^+^ CD4^+^ T_RM_ in the human lungs by confocal microscopy. The human lungs contain predominantly CD69^+^CD103^-^ CD4^+^ T_RM_ and 20-30% CD69^+^CD103^+^ CD4^+^ T_RM_, with frequencies varying among individuals^10,24^. Like CD103^+^CD8^+^ T_RM_, which are primarily embedded in the lung epithelium, CD103^+^CD4^+^ T_RM_ lined the airway epithelium, were distributed around airways, and did not form clusters (**Figure 1A**). On the other hand, CD69^+^CD4^+^ T_RM_ surrounded airways (**Figure 1B**), formed small and larger clusters in the lung parenchyma close to airways (**Figure 1C,D**). Interestingly, CD69^-^CD4^+^ T cells also appear to form small clusters in nearby airways (**Figure 1C**). These data show that CD103^+^CD4^+^ T_RM_ are distributed separately throughout the lung parenchyma and epithelium, whereas CD69^+^CD4^+^ T_RM_ and CD69^-^CD4^+^ T cells form clusters within the parenchyma and around airways.

**Figure 1:**
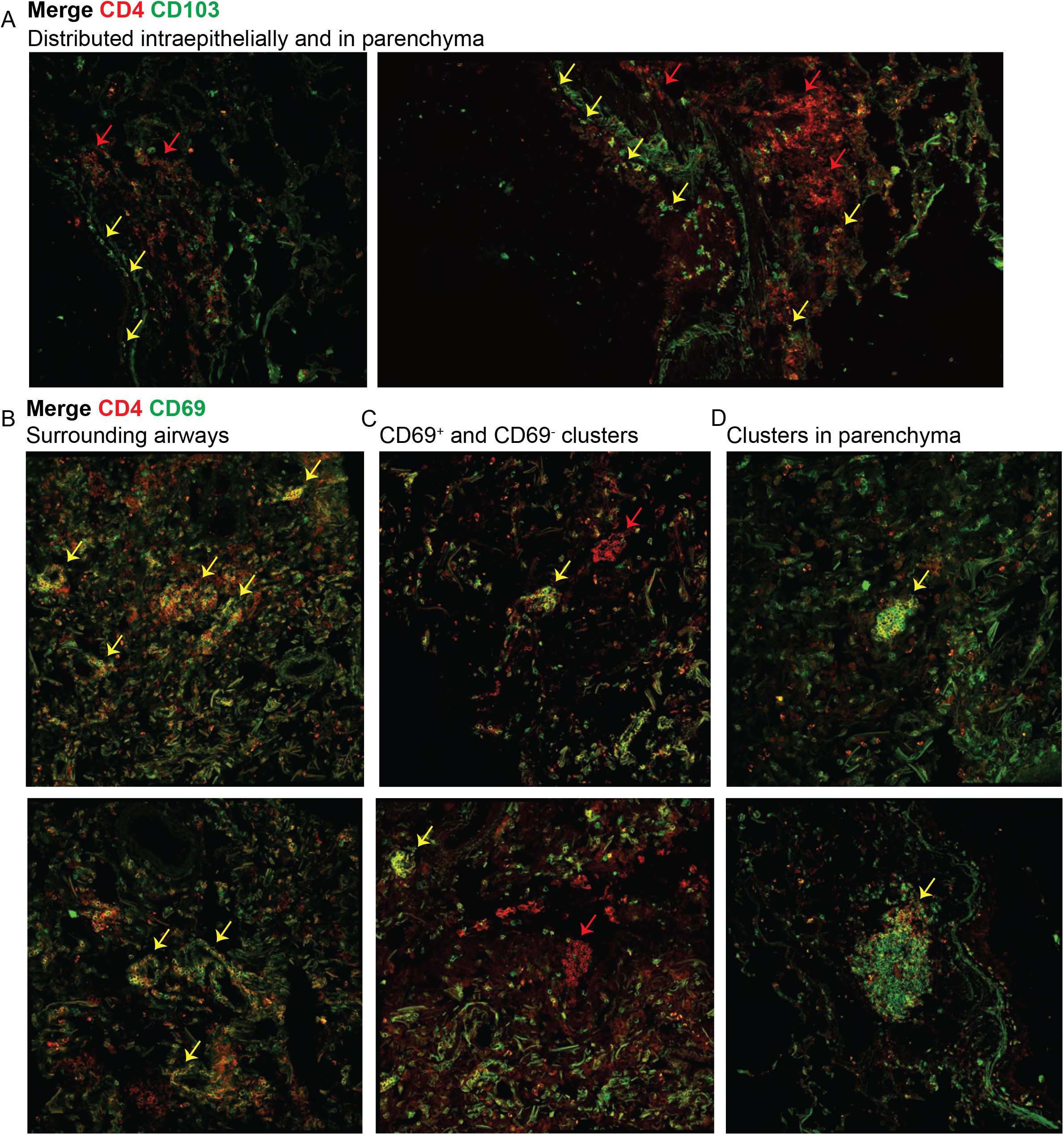
CD4^+^ T cells display distinct localization patterns in the human lungs. (**A**) Immunofluorescent staining for CD4 (red) and CD103 (green) demonstrating CD103^+^ (yellow) and CD103^-^ (red) CD4^+^ T cells in tissue-Tek embedded lung tissue at 40X magnification. (**B-D**) Immunofluorescent staining for CD4 (red) and CD69 (green) demonstrating CD69^+^ (yellow) and CD69^-^ (red) CD4^+^ T cells in various parts of the human lungs. Images are representative of N=2 from 2 independent experiments.

### Pathogen-specific CD4^+^ T_RM_ are enriched in the human lungs

To investigate whether the heterogeneous spatial organization also reflects functional diversity, we investigated CD4^+^ T_RM_ recognition of different airway pathogens, such as viruses, bacteria, and fungi. Therefore, we developed an *in vitro* stimulation assay to identify pathogen-specific CD4^+^ T_RM_ while preserving their original phenotypes. For standardization, we used identical antigen presenting cells (APCs) to stimulate the lung and blood-derived T cells. Paired PBMCs were labelled with cell trace violet (CTV) and the CD4^+^ and CD8^+^ cells were isolated from the lung mononuclear cells (LMCs). CD69 is often used as an activation marker, however, it is also used to identify T_RM_. To assess whether pathogen-specific CD4^+^ T cells in the lungs have a T_RM_ phenotype, the cells were stained with CD69 and excess CD69 antibody was washed from the medium before the start of stimulation. Thus, the cells expressing CD69 after the stimulation did not upregulate it during the assay, but already expressed it before activation. Labelled PBMCs and unlabeled CD4^+^/CD8^+^ lung cells were mixed and incubated overnight with different pathogen-derived peptides, proteins, or lysates (**Figure 2A**). Systemic and respiratory viruses, such as CMV, influenza, and RSV, as well as bacterial and fungal pathogens that colonize the airways, like *Aspergillus fumigatus, Staphylococcus aureus*, and *Streptococcus pneumoniae* were tested. The combined upregulation of CD40 ligand (CD40; CD154) and cytokine (TNF-α, IFN-γ, IL-2, IL-17a) production were used as the readout for antigen-specific stimulation. A sample was defined as reactive to a pathogen and called pathogen-specific when the frequency was at least two times higher than the unstimulated and at least 0.01% (general gating strategy **Supplementary figure 1A,B**).

**Figure 2:**
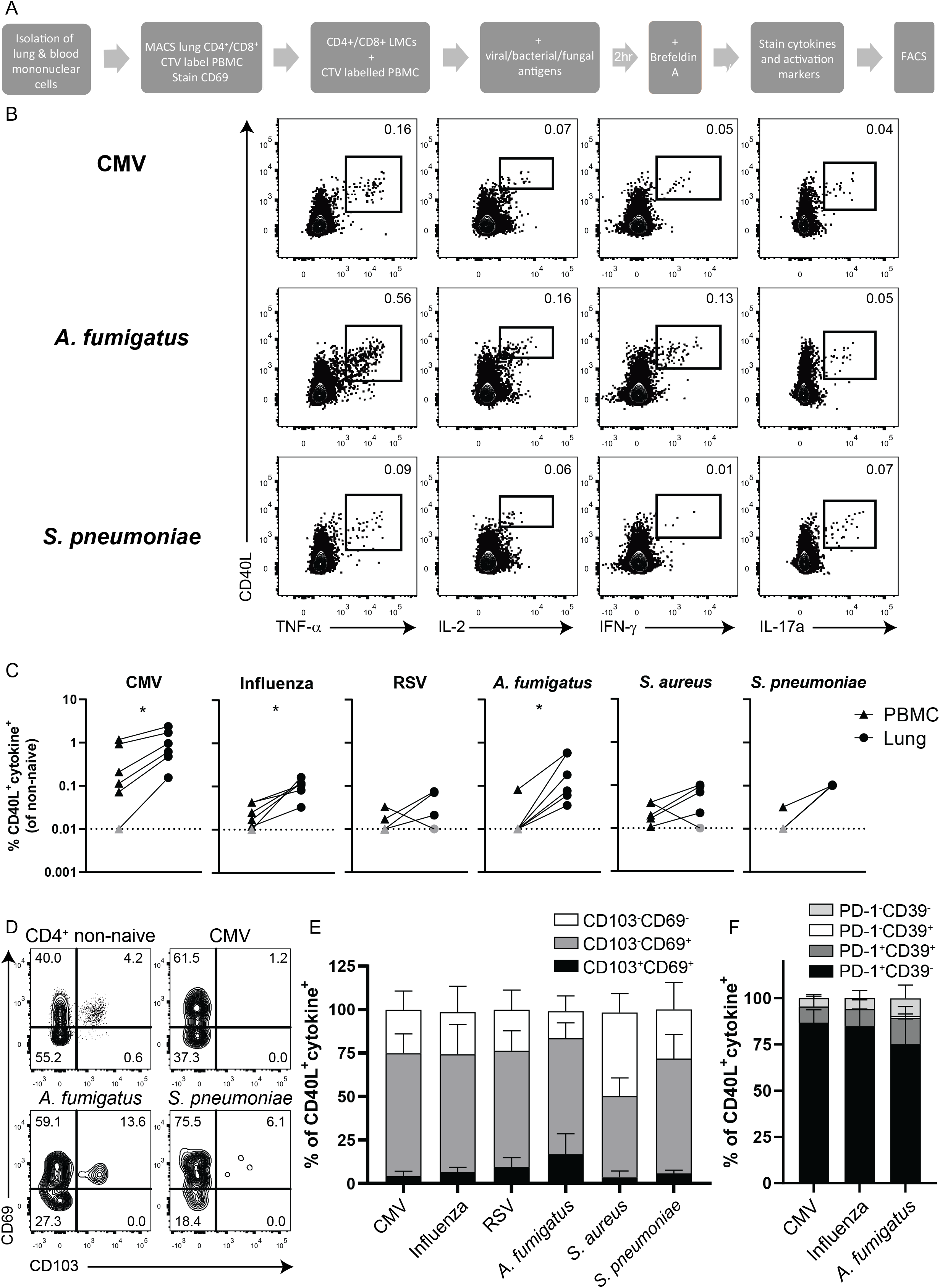
Pathogen-specific CD4^+^ T_RM_ are enriched in the human lungs. (**A**) The pathogen-specificity of CD4^+^ T cells in the lung and blood was determined using an *in vitro* stimulation assay. Briefly, lung mononuclear cells (LMCS) and PBMCs isolated from lung tissue and blood, respectively. MACS isolated CD4+/CD8+ LMCs and CTV-labelled PBMCs and stained with anti-CD69 prior to stimulation. CD4+/CD8+ LMCs and CTV-labelled PBMCs were mixed at a 1:5 ratio and incubated overnight with viral/bacterial/fungal antigens, with Brefeldin A added 2 hours into the incubation. (**B,C**) Frequencies of CMV-, influenza-, RSV-, *Aspergillus fumigatus-, Staphyloccocus aureus-*, and *Streptococcus pneumoniae*-specific blood (triangles) and lung (circles) non-naïve (NOT Boolean gating of naïve cells) CD4^+^ T cells were determined with flow cytometry by the upregulation of CD40L and cytokine (TNF-α, IL-2, IFN-γ, or IL-17a) production after peptide/protein/lysate stimulation, shown by representative contour plots for lung non-naïve CD4^+^ T cells (**B**) and quantified (**C**). (**D,E**) The expression of residency markers CD69 and CD103 was assessed on lung CMV-, influenza-, RSV-, *A. fumigatus-, S. aureus-*, and *S. pneumoniae*-specific CD4^+^ T cells, shown by representative contour plots (**D**) and quantified (**E**). (**F**) The expression of PD-1 and CD39 was determined on lung CMV-, influenza-, and *A. fumigatus*-specific CD4^+^ T cells. (**C**) Each symbol represents a unique sample and the line connects paired blood-lung samples. The significance was determined using Wilcoxon Signed Rank test. *p<0.05. N=6 for CMV, influenza, and *A. fumigatus*, N=4 for RSV, N=5 for *S. aureus*, and N=3 for *S. pneumoniae*. Grey symbols mark undetectable samples not reaching the detection limit (marked by dotted line at 0.01 on y-axis). The data is pooled from 7 independent experiments. (**E,F**) Stacked bars show the mean values with standard deviations. (**E**) N=6 for CMV, influenza, and *A. fumigatus*, N=4 for *S. aureus*, and N=3 for RSV, and *S. pneumoniae*. The data is pooled from 7 independent experiments. (**F**) N=3, data is pooled from 3 independent experiments.

Lung CD8^+^ T cells are enriched for recognition of respiratory viruses, but not systemic viruses^7^. On the contrary, the lungs were enriched for CMV-specific CD4^+^ T cells, as well as influenza-, and *A. fumigatus*-specific CD4^+^ T cells, with trends also observed for RSV, *S. aureus*, and *S. pneumoniae* (**Figure 2B,C**). In most patients, recognition of a pathogen in blood was predictive of recognition in the lung, with exceptions of *A. fumigatus-* and *S. pneumoniae*-specific CD4^+^ T cells. Interestingly, the phenotypes of the pathogen-specific CD4^+^ T cells differed between the two compartments, which may also point towards their spatial organization. In the lung, pathogen-specific CD4^+^ T cells mainly exhibited a CD103^-^CD69^+^ T_RM_ phenotype, whereas in blood these cells lacked surface expression of both CD103 and CD69 (**Figure 2D,E, Supplementary figure 2A,B**). Intriguingly, while influenza-specific lung CD8^+^ T cells are enriched in the CD103^+^ T_RM_ population^7^, very few CD4^+^ T cells specific for respiratory pathogens expressed CD103 in the lung. Approximately 20% of *A. fumigatus*-specific lung CD4^+^ T cells expressed both CD69 and CD103, but the majority were still CD103^-^CD69^+^. On the other hand, *S. aureus*-specific CD4^+^ T cells appeared to express less CD69 than other pathogen-specific cells. However, the phenotypes were not enriched compared to total non-naïve lung CD4^+^ T cells (**Supplementary figure 3C**). To investigate the activation status of the pathogen-specific CD4^+^ T_RM_, we determined the expression of PD-1 and CD39 on CMV-, influenza-, and *A. fumigatus*-specific CD4^+^ T cells. The expression of these two molecules is not altered by the stimulation, because the addition of Brefeldin A to the assay blocks the upregulation of surface proteins^25^ (**Supplementary figure 2D**). Therefore, we could assess which cells already expressed PD-1 and CD39 prior to the stimulation. Interestingly, the majority of the pathogen-specific CD4^+^ T cells expressed PD-1, but not CD39 (**Figure 2F, Supplementary figure 2E**). PD-1 is also a part of the T_RM_ core signature^8,10,11^. Therefore, the expression of CD69 and PD-1 and the lack of CD39 on the pathogen-specific CD4^+^ T cells strongly suggests that these cells are resident and not chronically activated or blood contaminants. Furthermore, the lack of CD103 expression indicates localization within clusters in the parenchyma rather than distribution along the lung epithelium.

### Human lung CD4^+^ T_RM_ exhibit distinct cytokine profiles corresponding to pathogen recognition

To investigate whether CD4^+^ T_RM_ also skew towards distinct Th1 or Th17 cytokine profiles, we examined the combined upregulation of CD40L and IFN-γ, IL-17a, and TNF-α production of lung CD4^+^ T_RM_ after overnight stimulation with polyclonal αCD3 mAb. In comparison to antigen-experienced CD4^+^ T cells in paired blood, antigen-experienced CD4^+^ T cells in the lungs produced significantly more IFN-γ, but similar amounts of IL-17a (**Supplementary figure 3A**). Lung CD4^+^ T cells produced IFN-γ and IL-17a, but these cytokines were not co-produced (**Figure 3A**). Furthermore, there were significantly more IFN-γ-producing Th1-type CD4^+^ T cells than IL-17a-producing Th17-type cells (**Figure 3B**). To ascertain whether these Th1 and Th17 cells exhibited a T_RM_ phenotype, the expression of CD69 and CD103 was measured on the cytokine-producing cells. The IL-17-producing cells were enriched for a CD69^+^CD103^-^ T_RM_ phenotype compared to both the total and IFN-γ-producing CD4^+^ T cells. Interestingly, contrary to CMV-, influenza-, and RSV-specific CD4^+^ T cells that were enriched for a CD69^+^CD103^-^ T_RM_ phenotype, the total IFN-γ producing CD4^+^ T cells skewed towards CD69^-^CD103^-^ phenotype (**Figure 3C-G**). This phenotypic enrichment was not observed for TNF-α producing cells, a cytokine not distinguishing specific helper types (**Supplementary figure 3B**). At the pathogen-specific level, the virus-specific CD4^+^ T_RM_ skewed towards IFN-γ production and were the most polyfunctional pathogen-specific CD4^+^ T cells, producing IFN-γ, TNF-α, and IL-2. *A. fumigatus*-specific CD4^+^ T_RM_ also produced more IFN-γ than IL-17a. A subset of CMV-specific lung CD4^+^ T cells produced IL-17a rather than IFN-γ. CD4^+^ T cells recognizing *S. aureus* and *S. pneumoniae* mainly produced IL-17a (**Figure 3H,I**). Altogether, these data demonstrate that the human lung contains both type 1 and 17 CD4^+^ T cells with distinct cytokine and phenotypic profiles allowing them to contribute to the adaptive immune response against a wide range of pathogens.

**Figure 3:**
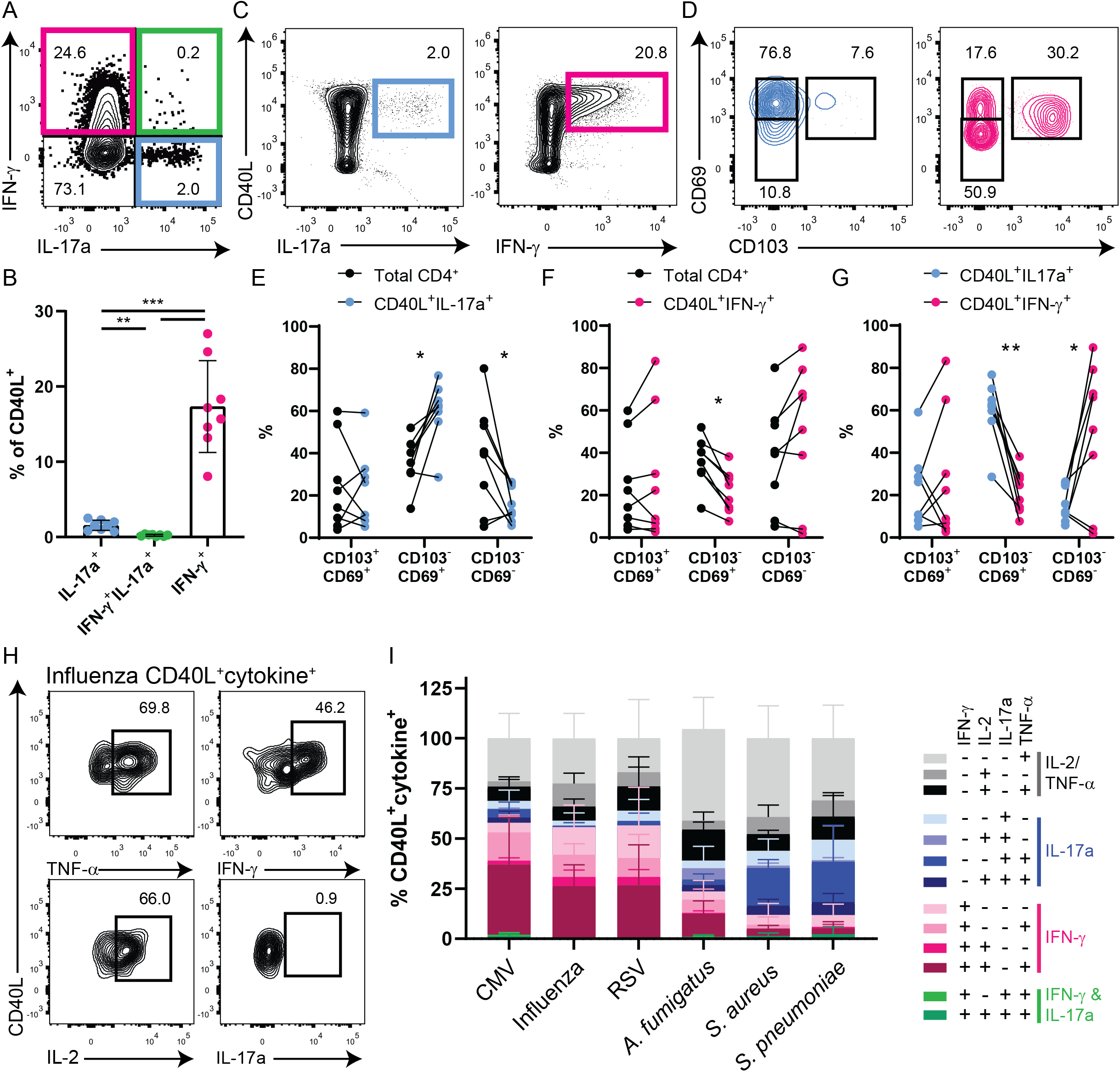
Lung pathogen-specific CD4^+^ T cells exhibit distinct cytokine profiles. (**A-B**) The production of IFN-γ (pink), IL-17a (blue), or both IFN-γ and IL-17a (green) by lung CD40L^+^CD4^+^ T cells was determined by flow cytometry after overnight αCD3 stimulation, shown by representative contour plot (**A**) and quantified (**B**). (**C,D**) The residency phenotype of CD40L^+^IL-17a^+^ and CD40L^+^IFN-γ^+^ (**C**) lung CD4^+^ T cells was measured by CD69 and CD103 expression (**D**), shown by representative contour plots. (**E-G**) The expression of CD69 and CD103 on the total CD4^+^ T cells was compared to CD40L^+^IL-17a^+^ (**E**) and CD40L^+^IFN-γ^+^ (**F**) CD4^+^ T cells and on CD40L^+^IL-17a^+^ compared to CD40L^+^IFN-γ^+^ CD4^+^ T cells (**G**). (**H,I**) The cytokine profiles of lung CMV-, influenza-, RSV-, *A. fumigatus-, S. aureus*-, and *S. pneumoniae*-specific CD4^+^ T cells were determined based on the production of TNF-α, IL-2, IFN-γ, and/or IL-17a, shown by representative contour plots (**H**) and quantified (**I**). (**B**) Each symbol represents a unique sample with the bar indicating the mean value and the significance was determined by one-way paired ANOVA with Tukey’s multiple comparisons test. (**E-G**) Each symbol represents a unique sample and the line connects paired samples. The significance was determined using two-way paired ANOVA with Sidak’s multiple comparisons test. *p<0.05, **p<0.01, ***p<0.001. (**B,E-G**) N=8 with data pooled from 8 independent experiments. (**I**) Stacked bars show the mean values with standard deviations. N=6 for CMV, influenza, and *A. fumigatus*, N=4 for *S. aureus*, and N=3 for RSV, and *S. pneumoniae*; the data is pooled from 7 independent experiments.

### Polyfunctional pathogen-specific bystander CD4^+^ T cells reside in tumors of NSCLC patients

Recently several research groups showed that many solid tumors contain pathogen-specific CD8^+^ T cells that infiltrate the malignant tissue. Therefore, we investigated whether TILs of NSCLC patients also consist of pathogen-specific CD4^+^ T cells, using the *in vitro* stimulation assay described above. Intriguingly, all tumor samples tested contained CD4^+^ T cells specific for the analyzed pathogens. Influenza-, RSV-, and *A. fumigatus*-specific CD4^+^ T cells were even enriched in the tumors compared to paired blood and comparable to frequencies detected in the lungs (**Figure 4A,B**). Responses against *S. pneumoniae* were not tested due to limited number of cells obtained from tumor samples. For the majority of samples and pathogens, presence of pathogen-specific CD4^+^ T cells in peripheral blood was predictive for these cells in the tumor. A large portion of the pathogen-specific, as well as total non-naïve, CD4^+^ TILs exhibited a residency phenotype, demonstrated by the expression of CD69 or CD69 and CD103, suggesting that these cells are not blood contaminants but reside in the tumors (**Figure 5A,B, Supplementary figure 2F**).

**Figure 4:**
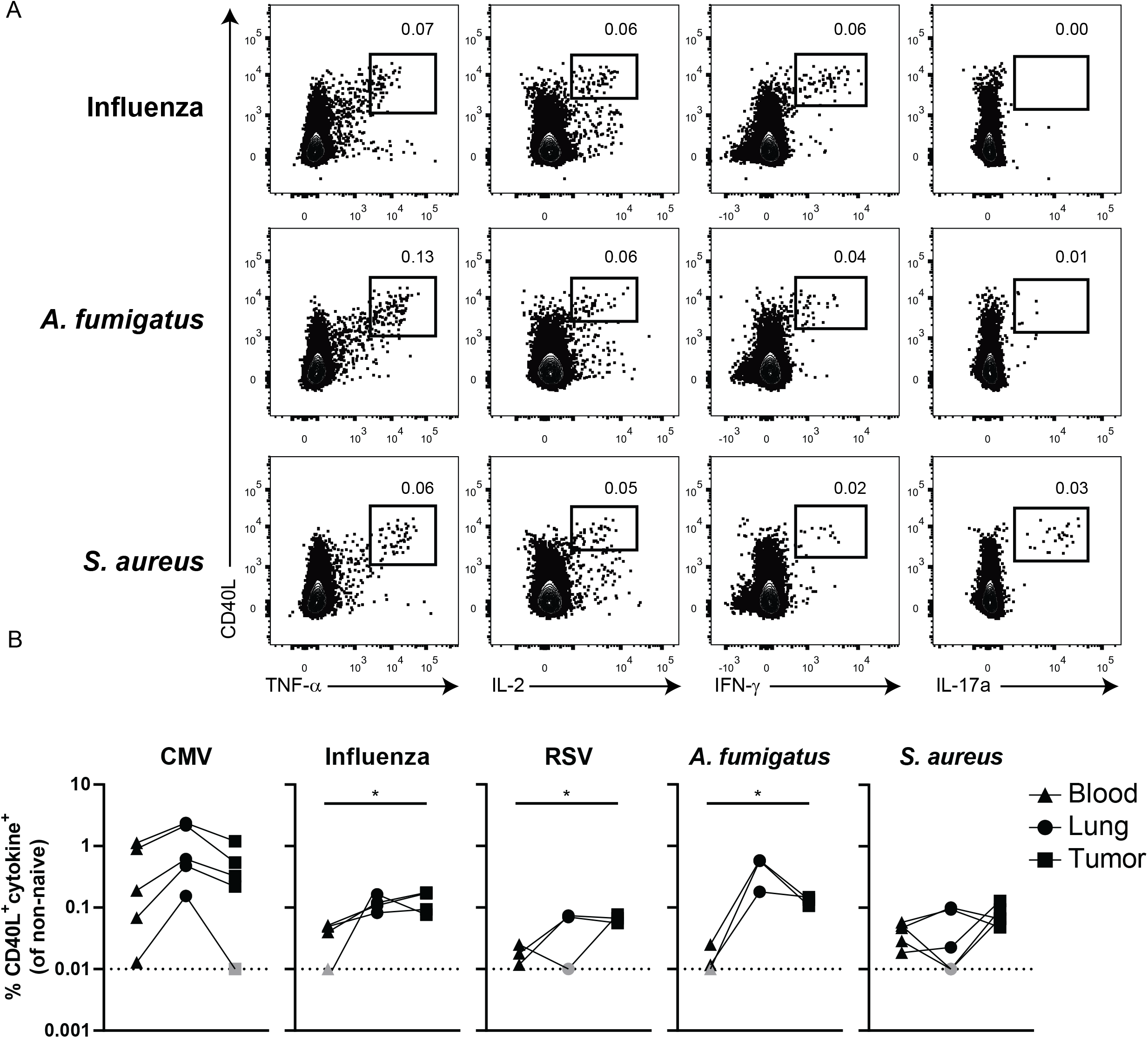
Pathogen-specific CD4^+^ T cells reside in tumors of NSCLC patients. (**A,B**) Frequencies of CMV-, influenza-, RSV-, *A. fumigatus-*, and *S. aureus*-specific blood (triangles), lung (circles), and tumor (squares) non-naïve CD4^+^ T cells were determined with flow cytometry by the upregulation of CD40L and cytokine (TNF-α, IL-2, IFN-γ, or IL-17a) production after peptide/protein/lysate stimulation, shown by representative contour plots for tumor non-naïve CD4^+^ T cells (**A**) and quantified (**B**). (**B**) Each symbol represents a unique sample and the line connects paired blood-lung-tumor samples. The significance was determined using paired one-way ANOVA with Tukey’s multiple comparisons test, *p<0.05. N=5 for CMV and *S. aureus*, N=4 for influenza, and N=3 for *A. fumigatus* and RSV. N differs for the pathogens due to limited number of cells isolated from some samples. The data is pooled from 7 independent experiments. Grey symbols mark undetectable samples not reaching the detection limit (marked by dotted line at 0.01 on y-axis).

**Figure 5:**
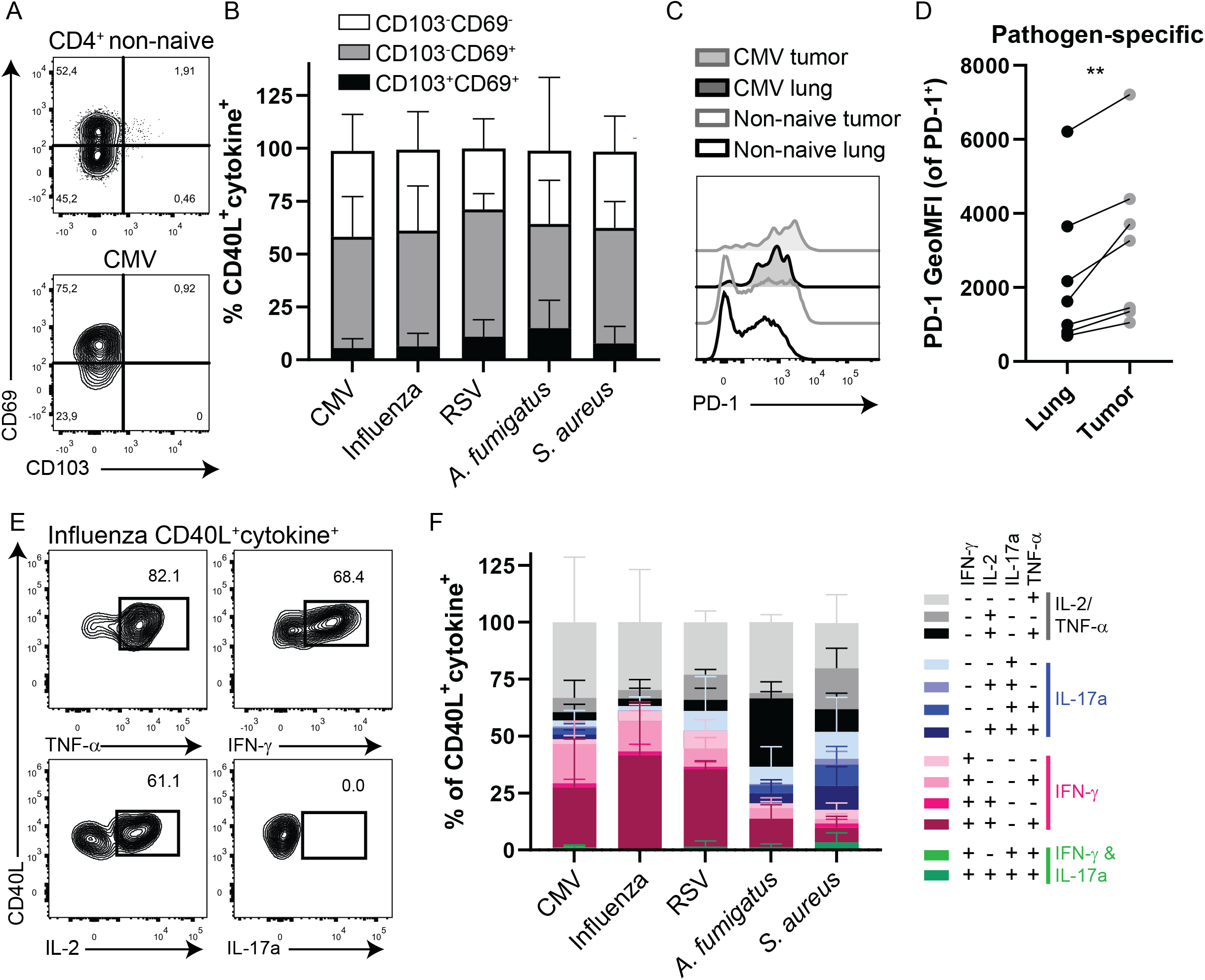
Tumor pathogen-specific CD4^+^ T cells exhibit a residency phenotype and retain polyfunctionality. (**A,B**) The expression of residency markers CD69 and CD103 was determined on tumor CMV-, influenza-, RSV-, *A. fumigatus-*, and *S. aureus*-specific CD4^+^ T cells, shown by representative contour plots (**A**) and quantified (**B**). (**C,D**) The expression of PD-1 was determined on lung and tumor pathogen-specific non-naïve CD4^+^ T cells, shown by representative half-off set histogram of tumor and lung CMV-specific and total non-naïve CD4^+^ T cells (**C**) and quantified as the geometric mean fluorescence intensity (GeoMFI) of PD-1 on PD-1^+^ cells (**E,F**) The cytokine profiles of tumor CMV-, influenza-, RSV-, *A. fumigatus-*, and *S. aureus*-specific CD4^+^ T cells were determined based on the production of TNF-α, IL-2, IFN-γ, and/or IL-17a, shown by representative contour plots (**E**) and quantified (**F**). (**B,F**) Stacked bars show the mean values with standard deviations. N=4 for CMV and influenza, N=3 for RSV and *A. fumigatus*, and N=5 for *S. aureus*; the data is pooled from 7 independent experiments. (**D**) N=7 paired pathogen-specific responses to CMV, influenza, *A. fumigatus*, and *S. aureus*, from 4 different patients; data is pooled from 4 independent experiments.

A major obstacle in anti-cancer immunity is the immunosuppressive tumor microenvironment (TME). Expression of inhibitory receptors, including PD-1, are often used to identify chronically stimulated T cells that lost the capacity to become activated and are therefore referred to as “exhausted”. To investigate whether PD-1 expression is altered on pathogen-specific CD4^+^ TILs, we compared PD-1 expression on pathogen-specific CD4^+^ T cells in paired lung and tumor samples. The majority of pathogen-specific CD4^+^ T cells expressed PD-1 regardless of the tissue (**Supplementary figure 2G**). However, the expression level of PD-1 was significantly higher on pathogen-specific CD4^+^ T cells in the tumor than in the paired lung samples, which indicates dysfunctionality (**Figure 5C,D**). Therefore, the cytokine profiles of the pathogen-specific CD4^+^ TILs was assessed. Intriguingly, the pathogen-specific CD4^+^ TILs displayed comparable polyfunctionality and pathogen-specific cytokine profiles as observed in the healthy lungs (**Figure 5E,F**). These data show that pathogen-specific CD4^+^ TILs in NSCLC resist exhaustion, and are licensed to respond with distinct effector functions, which may be therapeutically exploited for anti-tumor immunity.

## Discussion

Lung CD4^+^ T_RM_ serve an important role in the local immune response to airway pathogens. Localization is key to the functionality of T_RM_, as they strategically line the sites of pathogen entry, poised to trigger the alarm and curb the onset of infection. While transcriptional and phenotypic heterogeneity among CD4^+^ T_RM_ is well appreciated, the functional consequences, spatial organization and purpose of the different subsets remain unknown. Here, we demonstrate that in contrast to human lung CD8^+^ T_RM_ that line the airway epithelium, lung CD4^+^ T_RM_ also reside in the parenchyma surrounding airways and reside in clusters with other cells. While lung CD8^+^ T_RM_ are primarily involved in protection against respiratory viruses, CD4^+^ T_RM_ recognized a wide range of airway pathogens, including viruses, bacteria, and fungi. As such, it’s conceivable that CD4^+^ T_RM_ may also play a role in mediating the intricate balance between the lung microbiome and immune system. As different pathogens exploit different routes of entry, presumably the various pathogen-specific CD4^+^ T_RM_ reside in different niches requiring different phenotypic characteristics. Total IFN-γ producing lung CD4^+^ T cells skewed towards a CD69^-^ non-T_RM_ phenotype, compared to IL-17a producing CD4^+^ T cells enriched in the CD69^+^ T_RM_ population. We also showed that CD69^+^CD4^+^ T_RM_ and CD69^-^ CD4^+^ T cells form separate clusters within the lung parenchyma. This suggests differential clustering of Th1 and Th17 type CD4^+^ T cells in the parenchyma, presumably recognizing intracellular versus extracellular pathogens, respectively. However, the majority of CD4^+^ T cells specific for pathogens tested here, including viruses, expressed CD69. Perhaps the CD69^-^CD4^+^ T cells recognize pathogens not tested here or lack the molecules required for tissue retention, thereby enabling them to re-enter circulation and thereby escape the lung T cell repertoire.

The lifespan of T_RM_, especially in humans, is still not fully understood. In mice, CD8^+^ T_RM_ in the lungs diminish over time in the absence of antigen, whereas CD8^+^ T_RM_ persist in other tissues, such as the skin and gut^26-28^. Antigen persistence is required for prolonged replenishment and long-term maintenance of influenza-specific CD8^+^ T_RM_ in an influenza vaccination model^29^. On the other hand, in a human lung transplant study, donor lung CD8^+^ and CD4^+^ T_RM_ persist for at least 15 months post lung transplantation, demonstrating that human lung T_RM_ are maintained for at least a year^30^. Whether antigen is required for lung CD4^+^ T_RM_ maintenance remains under debate. Here, CMV-, influenza-, *A. fumigatus-*, and *S. pneumoniae*-specific CD4^+^ T_RM_ were enriched in the human lungs. CMV genomes are frequently detected in human lungs and CMV hides in alveolar macrophages^31,32^. Colonization of the upper respiratory tract by bacteria and fungi, such as *S. pneumoniae* and *A. fumigatus*, is common, especially in patients suffering from lung disease^33,34^. The *S. pneumoniae-* and *A. fumigatus*-specific CD4^+^ T_RM_ may maintain stable colonization by preventing bacterial outgrowth and infection, as is the case for Th17-type T_RM_ in the maintenance of *C. albicans* commensalism^35^. Conceivably, the persistence of CMV and possible colonization by bacteria and fungi facilitate the maintenance of the pathogen-specific CD4^+^ T_RM_ in the lungs. Microbiota-specific CD4^+^ T_RM_ are abundantly present in the guts of healthy individual and their function is altered during inflammation^36^. The symbiosis of the lung microbiota and CD4^+^ T_RM_ and their role in maintaining lung homeostasis and/or inducing inflammation, particularly in inflammatory lung disease, remain undetermined.

Pathogen-specific bystander CD4^+^ T cells with a residency phenotype also reside in tumors of NSCLC patients. In the last years, it has become evident that a large proportion of CD8^+^ and CD4^+^ TILs are not tumor-specific^21,37^. Furthermore, many patients fail to respond to immune checkpoint therapy or adoptive cell transfer regimens. Recently, an alternative approach was proposed. The use of anti-viral peptide therapy to trigger virus-specific bystander CD8^+^ TILs within tumors to activate surrounding innate and adaptive immune cells and recruit more immune cells to the tumor^23^. Thereby, transforming “cold” tumors into “hot” tumors. Viral-specific bystander CD4^+^ and CD8^+^ TILs produce cytokines, such as IFN-γ. While IFN-γ secreted by tumor-specific T cells exhibits direct cytopathic and cytostatic effects on tumor cells, IFN-γ can also diffuse throughout the tumor and indirectly alter tumor cells not recognized by T cells^38,39^. The generated cytokine field upregulates MHC class I expression on tumor cells making them more susceptible to tumor-specific CD8^+^ T cells. However, tumor cells also upregulate PD-L1 potentially increasing tumor immunosuppression. Instead of IFN-γ, fungi- and bacteria-specific CD4^+^ TILs produced IL-17a, which promotes tumor growth in several cancers, including lung cancer^40^. Here, we show that pathogen-specific bystander CD4^+^ TILs produce IFN-γ or IL-17a depending on the pathogen recognized. Therefore, while the use of pathogen-specific CD4^+^ T cells may add to this alternative approach, their potential anti-versus pro-tumor roles, especially in regard to IL-17a producing bystander CD4^+^ TILs, need careful assessment.

The major hurdle for current cancer immunotherapies is the immunosuppressive TME and resulting loss of T cell functionality. Here, we show that pathogen-specific CD4^+^ T cells in the tumor expressed higher levels of PD-1 than pathogen-specific CD4^+^ T cells in paired lung tissue, indicative of direct or indirect activation in the TME and consequently, dysfunctionality. On the contrary, pathogen-specific CD4^+^ TILs retained their polyfunctionality, suggesting that these cells are not chronically activated. CD4^+^ T cells, unlike CD8^+^ T cells, require antigen-presentation through MHC class II. However, many tumors are MHC class II negative^41^. Therefore, CD4^+^ T cells require local host APCs in tumors to present tumor antigens. CD4^+^ T cells can eliminate MHC class II negative tumor cells indirectly through IFN-γ production, presumably due to antigen presentation by host APCs and their consequent activation^42^. NSCLC tumors contain dendritic cells and B cells and CD4^+^ T cell clonal expansion correlates with the density of tumor B cells, suggesting that tumors contain local APCs able to present antigens to CD4^+^ T cells^43–45^. Therefore, the stimulation of polyfunctional pathogen-specific CD4^+^ TILs in tumors is possible.

Overall, CD4^+^ T_RM_ are distributed in distinct niches throughout the human lungs and recognize a multitude of pathogens. The spatial organization of the various pathogen-specific CD4^+^ T_RM_ remains to be elucidated. This may be important for vaccine design, specifically the route of vaccine administration, because it can severely affect the development of protective CD4^+^ T_RM_ and differs between pathogens, including *A. fumigatus*, *S. pneumoniae*, influenza, and coronaviruses^15,46–48^. Targeting CD4^+^ T_RM_ could be attractive for vaccine design as they are efficient multi-taskers, able to exert direct effector functions and guide the formation of optimal B cell responses and lung CD8^+^ T_RM_^49,50^. Furthermore, the presence of pathogen-specific CD4^+^ T cells in peripheral blood is a good predictor of the presence of these cells in tumors and could thus be used to screen patients for bystander specificity. Repurposing the polyfunctional pathogen-specific CD4^+^ TILs together with anti-viral CD8^+^ TILs detected in tumors of NSCLC patients provides an alternate avenue to boost tumor immunogenicity and efficacy of cancer immunotherapies.

## Materials and methods

### Subjects and study approval

Lung and tumor samples were obtained from non-small cell carcinoma (NSCLC) patients. The patients received a surgical resection of primary tumors as first line therapy without prior chemo- or radiotherapy. Blood was drawn from a central line at the start of surgery. The patients were recruited from Onze Lieve Vrouwe Gasthuis (OLVG), Amsterdam, the Netherlands. Written informed consent was given by all of the patients and donors before inclusion into the study. The Ethical ReviewBoard (ERB) of the METC/CCMO of the OLVG approved the study under the MEC-U number NL52453.100.15 according to the Declaration of Helsinki.

### Immunofluorescence

Lung samples were frozen in tissuetek and 8-10μM slices were stored at −80°C. Samples were fixed with dehydrated acetone and blocked with 1% BSA PBS. Sections were stained with AF594 anti-CD4 (Biolegend, clone RPA-T4), AF647 anti-CD69 (Biolegend, clone FN50) or AF647 anti-CD103 (Biolegend, clone Ber-ACT8). Nuclei were stained with Hoechst. SP8 (Leica) and ImageJ were used for imaging and analysis respectively.

### Isolation of peripheral blood, lung, and tumor mononuclear cells

Peripheral blood mononuclear cells (PBMCs) were isolated from heparinized blood samples using standard Ficoll-Paque density gradient centrifugation. Lung mononuclear cells (LMCs) and tumor mononuclear cells (TMCs) were isolated from the tissues as previously described^4,51^. In short, the tissue was cut into small pieces and incubated for 1 h at 37°C in digestion medium [RPMI with 20mM Hepes, 10% fetal calf serum (FCS), 50 U/ml DNAse type I (Sigma-Aldrich), 300 U/ml collagenase type 4 (Worthington)] on a roller. Before and after the digestion, the tissue was dissociated using gentleMACS Tissue Dissociator (Miltenyi). The digested tissue was passed through a flow-through chamber to achieve a single cell suspension. To isolate mononuclear cells from the cell suspension standard Percoll density gradient techniques were used. PBMC, LMC, and TMC samples were used directly for experimentation.

### *In vitro* stimulation assay for pathogen-specificity

For stimulation experiments with LMCs and TMCs, paired PBMC samples were first labelled with cell trace violet (CTV, Thermofisher) according to manufacturer’s instructions and CD4+ and CD8+ cells were isolated from LMCs using Miltenyi microbeads according to manufacturer’s protocol. Thereafter, CTV labeled PBMCs and CD4^+^/CD8^+^ LMCs and TMCs were stained with anti-CD69 BUV395 and washed prior to stimulation. 1-2 million paired CTV labelled PBMCs were mixed with 300,000 MACSed CD4^+^/CD8^+^ LMCs or TMCs in a 96-well plate. Cells were incubated with peptides, recombinant proteins, or lysates at 37°C (5% CO2). Activation with soluble mAb αCD3 (Thermofisher, clone HIT3A) was used as a positive control and for the negative control, cells were incubated overnight in medium, referred to as unstimulated cells. After two hours, Brefeldin A (Thermofisher) was added and the cells were further incubated overnight.

### Peptides, recombinant proteins, and lysates

The following peptides pools were used at a final concentration of 100ng/ml; PepTivator Influenza A H1N1 (NP, MP1, and MP2; Miltenyi Biotech), and pepMix RSV (Glycoprotein G, and fusion protein; JPT Peptide Technologies). Recombinant CMV pp65 protein was used at a final concentration of 10 μg/mL (Miltenyi Biotech, Swiss-Prot P06725). *S. aureus* (NCTC 8325; 5×10^9^ CFU/ml), *S. pneumoniae* (TIGR4) and *A. fumigatus* (Miltenyi Biotech; 1mg/ml) lysates were diluted 1:1000 for the assays. Heat inactivated *S. aureus* and *S. pneumoniae* bacterial lysates were provided by B. Bardoel and S. Rooijakkers.

### Flow cytometric analysis

Cells were stained according to the manufacturer’s instructions. Briefly, cells were stained with anti-CD69 BUV395 (BD, clone FN50) on the surface and washed prior to stimulation. After the stimulation, cells were surface stained with anti-CD103 BV711 (Biolegend, Ber-ACT8), anti-CD45RA BUV563 (BD, clone HI100), anti-CD27 BV650 (Biolegend, clone 0323), anti-PD-1 PE-Cy7 (BD, clone EH12.2H7), and anti-CD39 BV510 (Biolegend, clone A1). Cells were then fixed and permeabilized with the Foxp3 Staining Buffer Set (Thermofisher), followed by intracellular staining of the cells with anti-CD3 BV605 (eBioscience, clone OKT3) or BUV661 (BD, clone UCHT1), anti-CD4 BUV737 (BD, clone SK3), anti-CD8 BUV805 (BD, clone SK1), anti-CD40L PeDazzle594 (Biolegend, clone 24-31), anti-CD137 AF647 or PeCy7 (Thermofisher Scientific, clone 4B4-1), anti-TNF PeCy7 or FITC (BD, Mab11), anti-IFNγ eF450 or BV785 or Pe (Biolegend, clone 4S.B3), anti-IL17a BV605 (Biolegend, clone BL168), and anti-IL-2 BB700 or APC (BD, clone MQ1-17H12). Samples were analyzed in 0.5% FCS PBS and acquired on a BD FACSymphony. Data analysis was performed using FlowJo (TreeStar, Version 10.0.7).

### Analysis of pathogen-reactivity

NOT Boolean gating of naïve CD4^+^ T cells was used to gate on non-naïve CD4^+^ T cells to normalize for the abundance of naïve cells in peripheral blood and the lack of these cells in the lungs. Boolean OR combination gating was performed in FlowJo to also calculate the frequencies pathogen-specific non-naïve CD4^+^ T cells based on upregulation of CD40L and production of IFN-γ, TNF-α, IL-2, and/or IL-17a; defined as the percentage of cells upregulating CD40L and producing at least one cytokine. To distinguish between responders and non-responders, a detected response was deemed positive if it was at least two times the magnitude of the unstimulated condition and at least 0.01%. Only these responses were used for further analysis.

### Statistics

The significance of the results was calculated with GraphPad Prism (version 8.0.2) using 2-way ANOVA with Sidak’s multiple comparisons test, RM 1-way ANOVA with Tukey’s multiple comparisons test or Wilcoxon’s Signed Rank test as indicated per figure legend.

## Data availability

The datasets generated during and/or analyzed during the current study are available from the corresponding author on reasonable request.

## Acknowledgements

We would like to thank all patients who participated in this study. Furthermore, we’d like to thank all the study nurses and doctors who participated in the study. We would also like to extend our gratitude to Erik Mul, Simon Tol, and Mark Hoogenboezem of the Sanquin Central Facility for their assistance. We would also like to thank Regina Stark for carefully reading the manuscript. This study was funded by Sanquin Blood Supply project grant PPOC, Jochem van Loghem foundation and the Dutch Research Council (NWO) Veni grant.

## Author contributions

AO and PH designed the study. AO, FM, RH, CG performed the experiments in this manuscript. AO performed the data analysis. All authors contributed to interpretation and discussion of data. AO and PH wrote the manuscript. All authors read and approved the manuscript.

