## Supplementary figures for "Polyfunctional pathogen-specific CD4^+^ T cells reside in the lungs and tumors of NSCLC patients"

Supplementary figure 1: General gating strategies used in the manuscript.
(A) The general gating strategy used throughout the manuscript, starting with lymphycytes and single cells, follwed by live (NearIR$^{-}$) CTV+ PBMCs and CTV- LMCs, CD3+ T cells, gating out aggregates, CD4+, gating on naive cells and using Boolean NOT gating to have non-naive (antigen-experienced) cells. (B) Representative contour plots showing expression of CD40L and cytokine (TNF-a, IL-2, IFN-g, and IL-17a) production by unstimulated lung (top plots), unstimulated PBMC (middle plots) and CMV-stimulated PBMC (bottom plots) non-naive CD4$^{+}$ T cells.

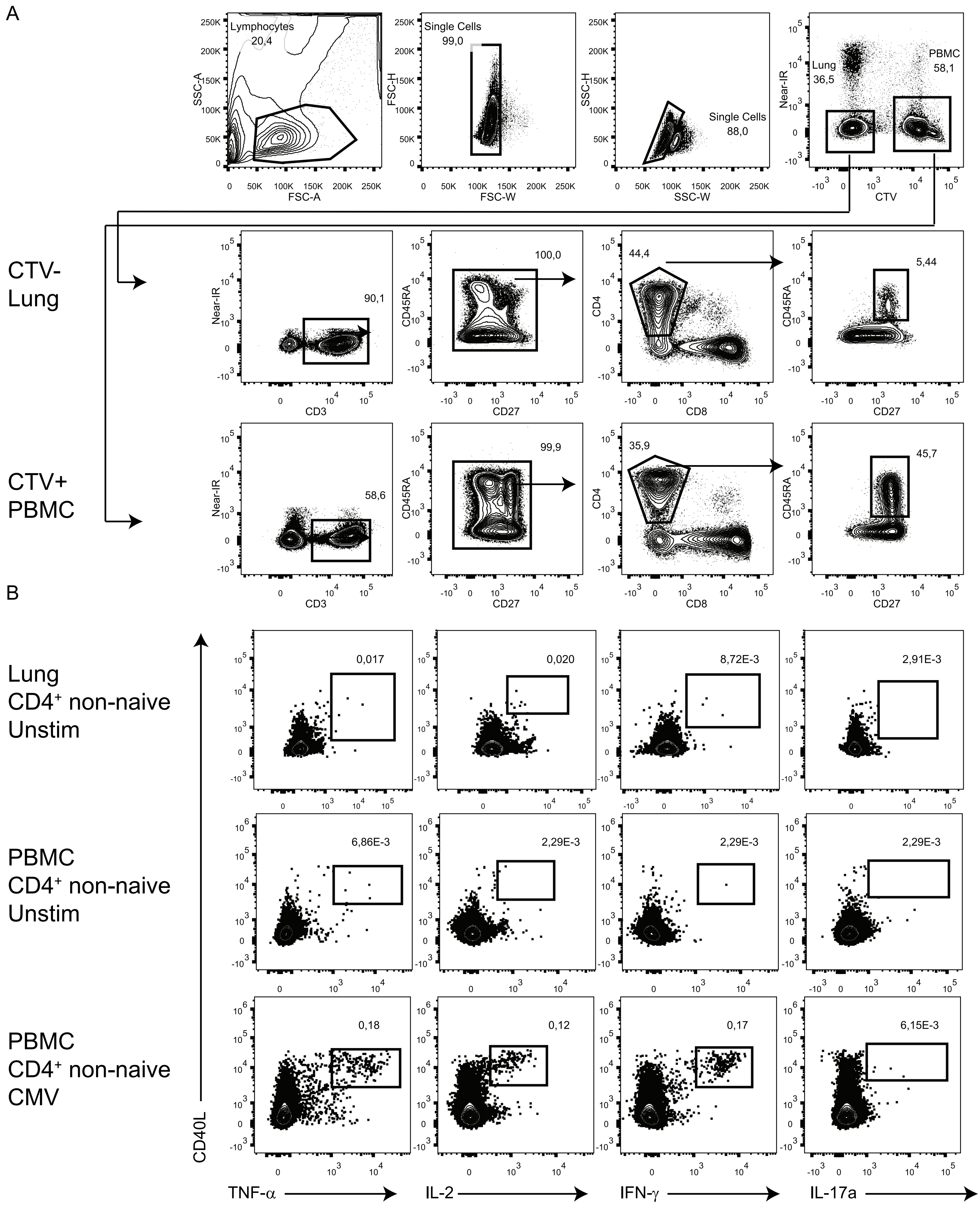

Supplementary figure 2: The phenotype of pathogen-specific $CD4^+$ T cells.
(A,B) The expression of CD69 and CD103 was determined on $CD40L^+cytokine^+$ $CD4^+$ T cells, shown by representative controur plots (A) for lung influenza, RSV, and *S. pneumoniae* stimulated, for PBMC CMV, influenza, and *S. aureus* stimulated, and quantifed for PBMCs (B). (C) The expression of CD69 and CD103 was determined on total non-naive lung $CD4^+$ T cells and quantified. (D,E) The expression of CD39 and PD-1 was measured on total unstimulated and αCD3-stimulated non-naive lung $CD4^+$ T cells (D) and $CD40L^+$ $cytokine^+$ CMV, influenza, and A. fumigatus stimulated lung $CD4^+$ T cells (E). (F) The expression of CD69 and CD103 was determined on total non-naive tumor $CD4^+$ T cells and quantified. (G) The frequency of $PD\text{-}1^+$ cells of pathogen-specific lung and tumor $CD4^+$ T cells was quantifed. (B,C,F) The bars show mean values with standard deviations, (B) N=5 for CMV, influenza, and *S. aureus* and N=3 for RSV, (C) N=7, (F) N=7; data is pooled from 7 independent experiments. (G) Each symbol represents a unique sample and the line connects paired samples. N=7, with data pooled from 3 independent experiments.

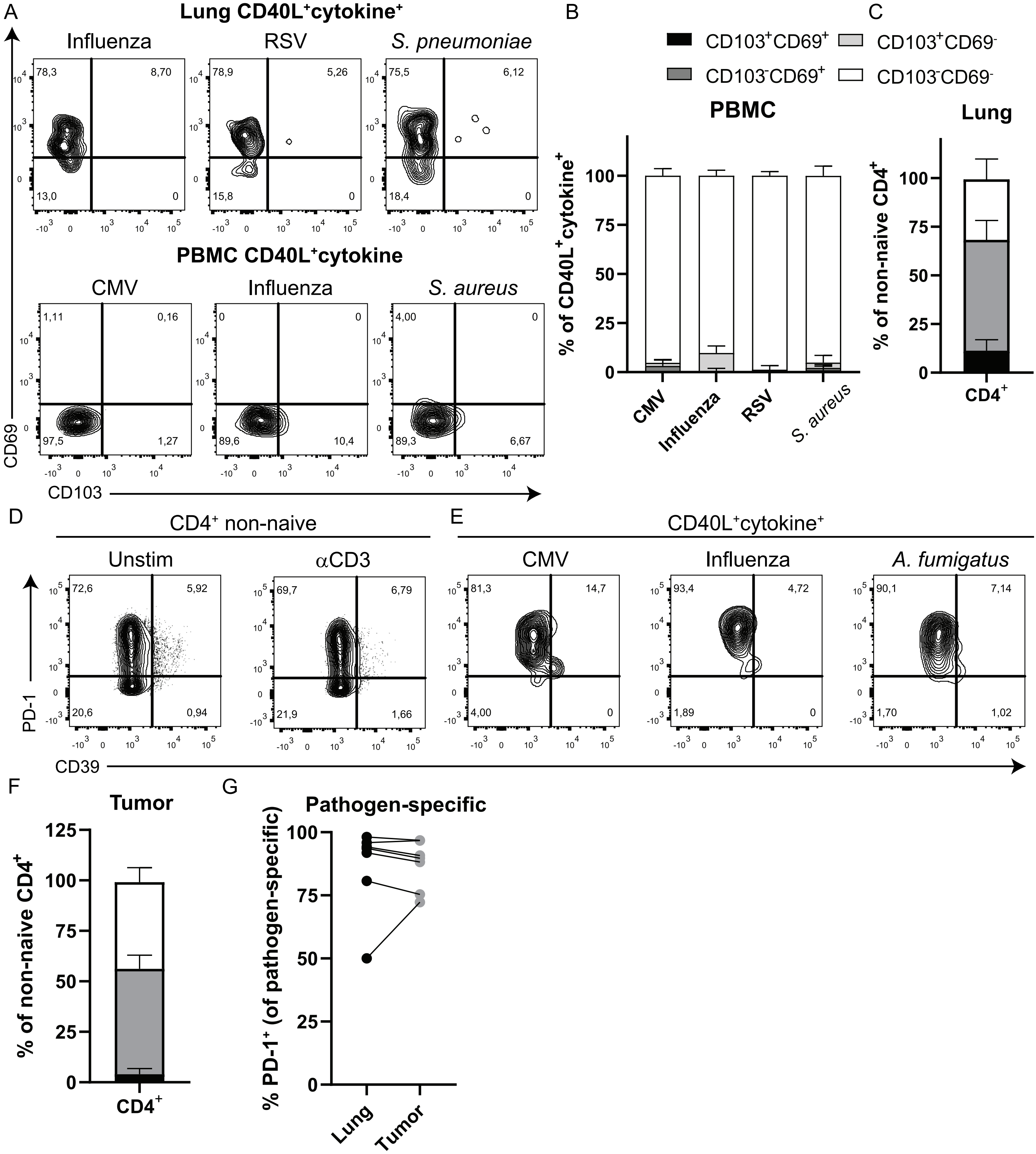

Supplementary figure 3: Cytokine profiles of lung CD4⁺ T cells.
(A) Production of IFN-γ, IFN-γ and IL-17a, or IL-17a by CD40L⁺CD4⁺ T cells from lung (circles) and blood (triangles) determined by αCD3 stimulation. (B) Residency phenotype of lung CD40L⁺TNF-α⁺CD4⁺ T cells versus total CD4⁺ T cells determined by CD69 and CD103 expression.

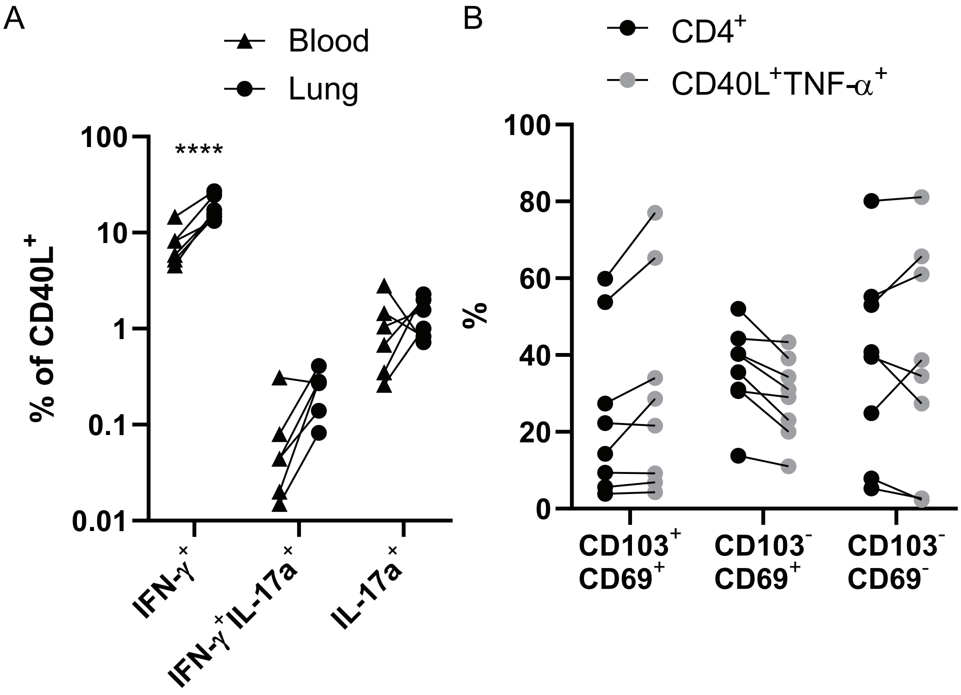
